# Accurate eQTL prioritization with an ensemble-based framework

**DOI:** 10.1101/069757

**Authors:** Haoyang Zeng, Matthew D. Edwards, Yuchun Guo, David K. Gifford

## Abstract

Expression quantitative trait loci (eQTL) analysis links sequence variants with gene expression change and serves as a successful approach to fine-map variants causal for complex traits and understand their pathogenesis. In this work, we present an ensemble-based computational framework, EnsembleExpr, for eQTL prioritization. When trained on data from massively parallel reporter assays (MPRA), EnsembleExpr accurately predicts reporter expression levels from DNA sequence and identifies sequence variants that exhibit significant allele-specific reporter expression. This framework achieved the best performance in the “eQTL-causal SNPs” open challenge in the Fourth Critical Assessment of Genome Interpretation (CAGI 4). We envision EnsembleExpr to be a powerful resource for interpreting non-coding regulatory variants and prioritizing disease-associated mutations for downstream validation.

## 1 Introduction

Genome-wide association studies (GWAS) have identified thousands of variants relevant to complex traits or diseases (McCarthy *et al.*, 2008; Manolio, 2010; Stranger *et al.*, 2011; Hindorff *et al.*, 2009). However, as most of these variants reside in non-coding regions of the genome (Hindorff *et al.*, 2009; Frazer *et al.*, 2009), distinguishing the causal variants from the ones in strong linkage disequilibrium (LD) remains challenging. Expression quantitative trait loci (eQTL) analysis has been widely used to assist in fine-mapping the causal mutations and provide immediate insight into their biological basis (Cookson *et al.*, 2009). But meanwhile, similar to GWAS, the statistical power of eQTL analysis is constrained by the complicated LD structure of the human genome and the statistical multiple-comparison burden from the large number of variant-gene pairs to investigate.

The massively parallel reporter assay (MPRA) is an efficient way to systematically dissect transcriptional regulatory elements (Melnikov *et al.*, 2012). In MPRA, a large number of synthesized DNA elements and corresponding sequence tags are cloned into plasmids to form reporter constructs and are transferred to cells. The expression of the tag is subsequently assayed by high-throughput sequencing. Tewhey et al. further improved the efficiency and reproducibility of MPRA (Tewhey *et al.*, 2016) to interrogate the expression level of reference and alternate alleles of 9,116 variants linked to 3,157 eQTLs. With this dataset, they discovered hundreds of variants with significantly different expression between the two alleles (allele-specific expression). In the Fourth Critical Assessment of Genome Interpretation (CAGI 4), this dataset was used as the training and test sets in the “eQTL-causal SNPs” challenge to identify the best computational approaches to predict (reporter) expression level from DNA sequence and to classify which sequence variants will lead to allele-specific expression.

In this work, we present a computational framework, EnsembleExpr, that outperformed all the competing methods in both parts of the challenge. The performance of EnsembleExpr is robust to various evaluation metrics. As an ensemble model, EnsembleExpr achieves performance superior to any single component by integrating complementary features of different sources and properties. We also demonstrate how a sufficient range of sequence-based annotation of functional elements is crucial to achieving accurate prediction of gene expression levels.

## 2 Background

### 2.1 Datasets in CAGI4 eQTL challenge

Tewhey et al. (Tewhey *et al.*, 2016) identified all the variants (range = 1 to 205, mean = 2.87, median = 1) in perfect LD with 3,157 eQTLs drawn from the Geuvadis RNA-seq dataset of lymphoblastoid cell lines (LCLs) from individuals of European ancestry (Consortium *et al.*, 2012; Lappalainen *et al.*, 2013). For each variant, the 150-bp flanking sequence of each of the two alleles were synthesized with the corresponding allele centered at the middle of the synthesized oligonucleotide. With these sequences as the library, MPRA experiments were carried out in two lymphoblastoid cell lines.

These data were split into three groups of similar sizes. The first group is the training set. It consists of 3,044 variants, in which the normalized plasmid counts, RNA counts, log2 fold expression level (“**Log2FC**”), expression p-value, multiple-testing corrected p-value and whether the expression for either of the two alleles is significantly high (Regulatory Hit or “**RegHit**”). For each variant, the dataset also includes the log2 ratio (alternative/reference) of expression (“**LogSkew**”), allelic skew p-value, allelic skew FDR and whether the change in expression is significantly large (“**emVar**”).

The next two groups were used as the test set, the labels of which were not provided to the participants in the challenge. The second group consists of 3,006 variants, and the participants were required to submit allele-specific expression predictions (“Log2FC”) and whether it is significant (“RegHit”). The last group consists of 3,066 variants, 401 of which have at least one allele with strong expression (“RegHit”). For these 401 variants, the participants were asked to predict allelic change of expression (“LogSkew”) and whether it is significant (“emVar”). For all of the three groups, only the genomic location and the sequences of the alleles were provided.

### 2.2 Tasks in CAGI4 eQTL challenge

#### Expression Prediction

In this task, the participants needed to submit predictions and confidence estimates for the expression level (“Log2FC”, real value) and whether the expression is significant (“RegHit”, binary label) for the second group of data.

#### Allele-specific Expression Prediction

In this task, the participants needed to submit predictions and confidence estimates for the change of expression between two alleles of a variant (“LogSkew”, real value) and whether the change is significant (“emVar”, binary label) for the third group of data.

## 3 Methods

### 3.1 Features

Sequence-based features were generated for the candidate regulatory regions given in the challenge (Figure 1A). First, 150-bp probe sequences were obtained for both studied alleles, as described in the challenge input files. Then for the set of sequences, we applied several computational approaches, including Kmer-Set Motif (KSM, in preparation), DeepSEA (Zhou and Troyanskaya, 2015), DeepBind (Alipanahi *et al.*, 2015) and ChromHMM (Ernst and Kellis, 2012) to derive sets of functional features that we hoped would help us predict expression levels.

**Figure 1:**
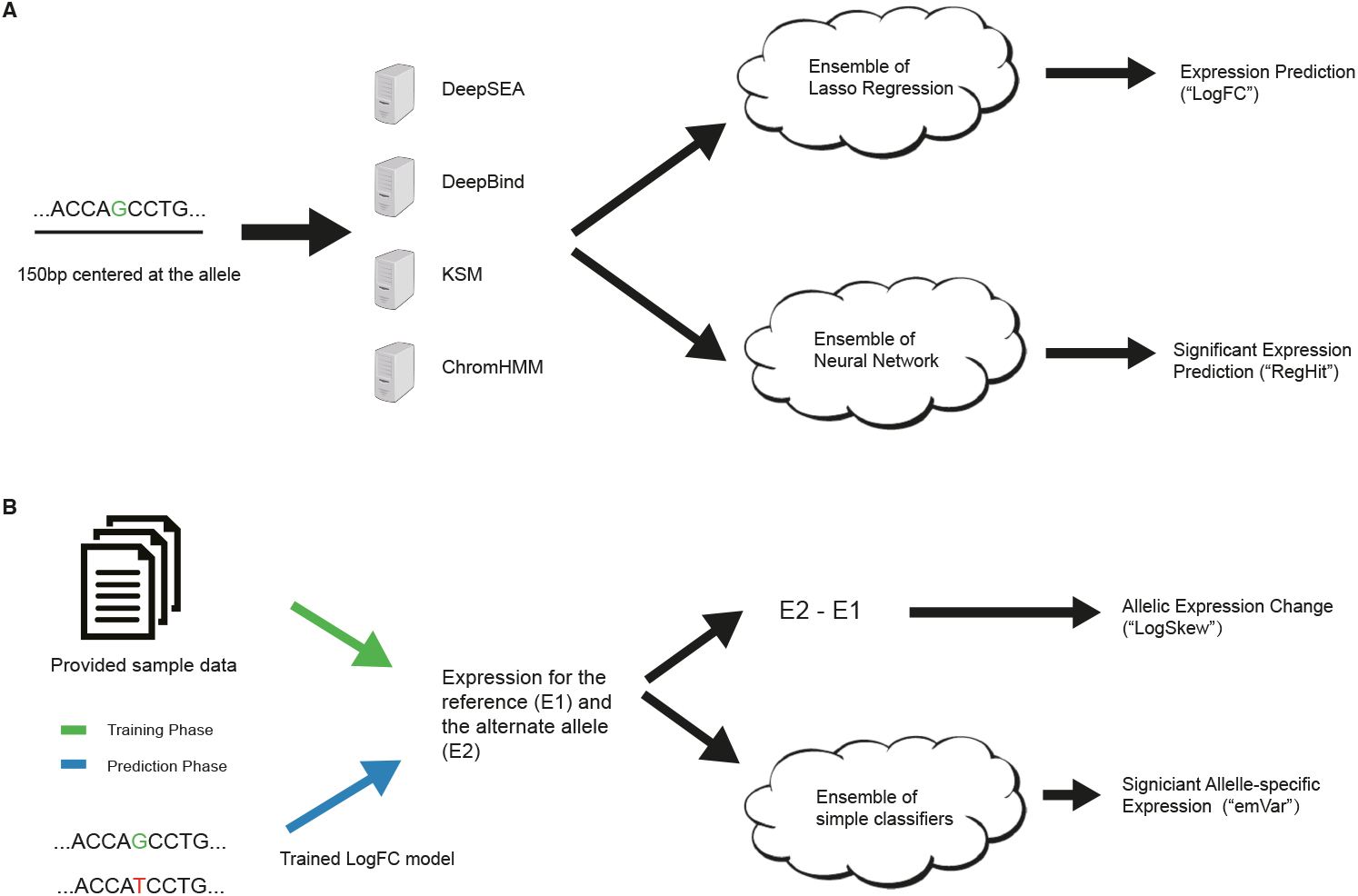
The schematic of EnsembleExpr. (A) The 150-bp sequence centered at the queried allele is taken as input to four computational models to generate functional features, on which two ensemble models are built to make expression predictions (“log2FC”) and significance estimates (“RegHit”). (B) In the training phase, the provided expression levels of the two alleles of each variant are used to train an ensemble model of significant allele-specific expression (ASE). In the testing phase, we first apply the trained expression model in (A) to generate expression predictions, which are then given to the significant ASE model to make predictions (“emVar”). The difference of the predicted expression levels is directly output as the prediction for allelic expression change (“LogSkew”).

Specifically, we used the DeepSEA probabilistic model to analyze the 150-bp sequences and obtain allele-specific signal predictions for 919 DNase-seq, transcription factor ChIP-seq, and histone mark ChIP-seq experiments. Similarly, we applied the DeepBind model to the same sequences and generated allele-specific predictions for the binding affinities of 538 distinct transcription factors. In addition, a KSM model trained on 57 ENCODE ChIP-seq experiments for a lymphoblastoid cell line (GM12878) was used to produce predictions for transcription factor binding affinities. Chromatin state annotations from the NIH Roadmap Epigenomics (Kundaje *et al.*, 2015) project were also compiled for all regions and used as one-hot encoded binary features.

### 3.2 Computational model

#### Expression Prediction Task

Armed with this set of potentially predictive features for expression levels, many of which are allele-specific, we used an ensemble of regularized regression and classification models to predict allele-specific expression values and regulatory hit status based on the provided training data (Figure 1A).

Specifically, we trained multiple LASSO regression models to predict the log (normalized) expression levels for each allele using the DeepSEA features alone, the DeepBind features alone, DeepSEA and KSM features combined, and DeepSEA along with KSM and chromatin state annotations. All learning algorithms were tuned by cross-validation, and the various feature sets were chosen using a heuristic manual analysis. We averaged the LASSO model predictions to produce the final predictions and took the standard deviation of the separate predictions as confidence estimates. For the binary prediction task (“RegHit”), we trained a one-layer neural network with 400 neurons on the same four sets of features described previously.

#### Allele-specific Expression Prediction Task

For allelic expression change (“LogSkew”) prediction, given that “LogSkew” is defined and calculated as the expression difference between the two alleles, we decided to directly utilize the “Log2FC” expression model we trained in the previous section instead of training a new model. Therefore we applied the trained “Log2FC” model to generate expression predictions for each allele in the held-out test set to submit. Then for each variant, we took the difference in predicted expression levels between the reference and the alternate alleles as our “LogSkew” prediction (Figure 1B).

For predicting “emVar” labels (allele-specific expression status), we trained on the actual allelespecific expression levels provided in the sample data. An ensemble of binary classification models was considered, with all regularization parameters tuned by cross-validation (Figure 1B). Models used in the final ensemble included linear regularized logistic regression, kernel regularized logistic regression, k-nearest neighbors, support vector machine (SVM) with linear kernel and SVM with radial basis function kernel. The predictions of all models were combined to form the final probability estimate along with a measure of confidence in the prediction. After training, we first ran our prediction module in the previous task (Expression Prediction) to generate predictions of allele-specific expression, on which we applied the trained model here to make predictions of significant allele-specific expression (“emVar” hits) for the held-out challenge dataset.

## 4 Results

### 4.1 EnsembleExpr outperforms competing approaches in CAGI eQTL challenge

We assessed the prediction from EnsembleExpr and other competing methods in the challenge. For predicting log (normalized) expression levels and expression levels between two alleles (“LogSkew”), both of which are regression tasks, we used Spearman’s rank correlation coefficient which is non-parametric and stable with the scale of the values. For predicting significant expression (“RegHit”) and significant allele-specific expression (“emVar”), both of which are binary classification tasks, we chose two benchmarks: receiver operating characteristic (ROC) and precision recall curve (PRC). ROC evaluates how the true positive rate changes with the false positive rate, where a random prediction would be along the diagonal with an area under curve (AUC) of 0.5 and a better model would have larger AUC. PRC shows how the precision changes with increasing recall (true positives), where the desired model should maintain high precision for large recall.

EnsembleExpr outperformed all the competing methods in both tasks. In the first part of the challenge, expression predictions from EnsembleExpr correlate the best with the experimental observations (Table 1, a Spearman correlation of 0.485 for the reference allele and 0.470 for the alternate allele). In predicting significant expression (“RegHit”), EnsembleExpr is the only model with an auROC > 0.8 and an auPRC > 0.5 (Table 1, Figure 2A).

**Figure 2:**
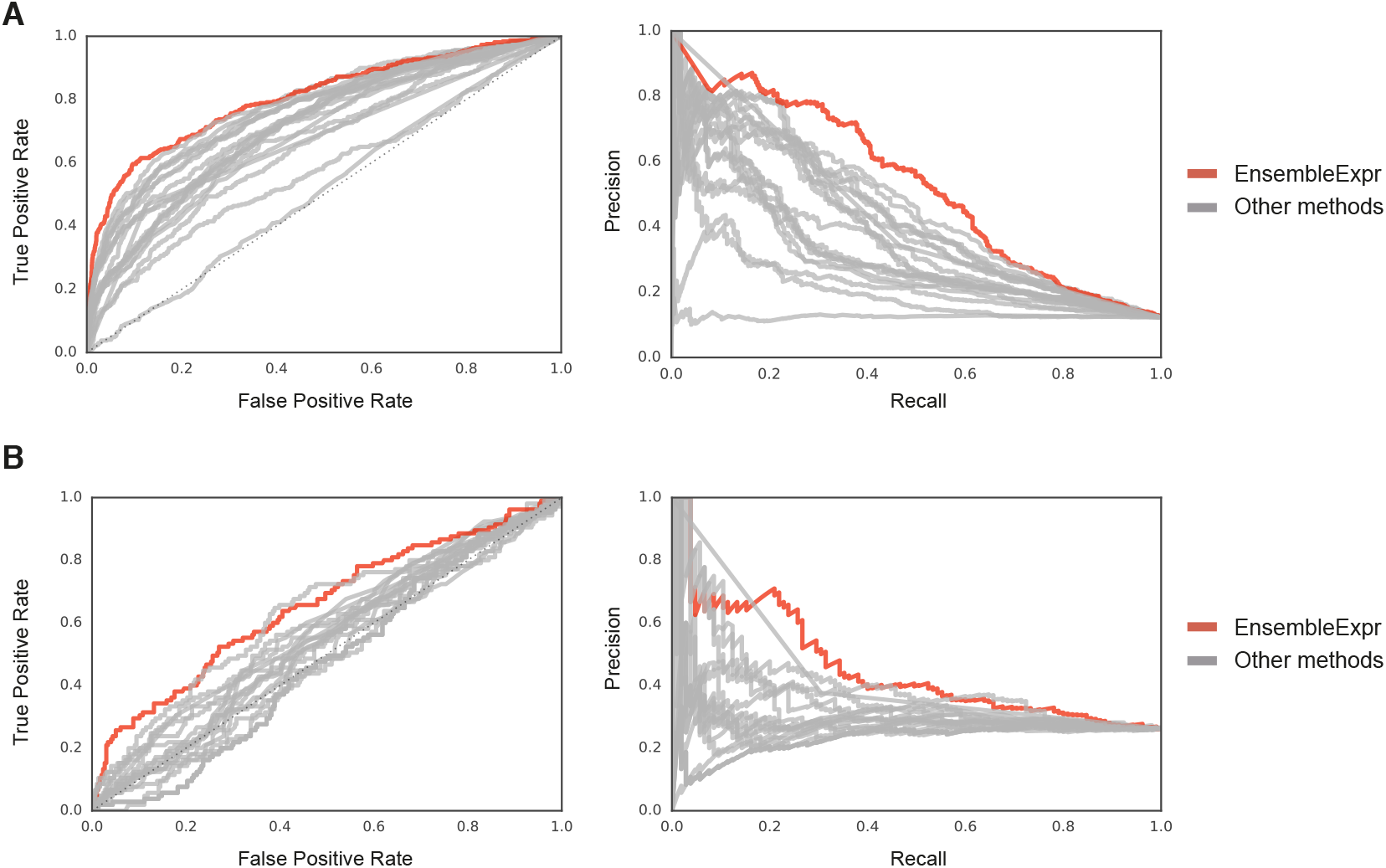
EnsembleExpr outperformed all the competing methods. (A) The area under ROC (auROC, left) and area under precision-recall curve (auPRC, right) for EnsembleExpr (red) and other methods (grey) in predicting significant expression. (B) The auROC (left) and auPRC (right) for EnsembleExpr (red) and other methods (grey) in predicting significant allele-specific expression.

**Table 1:** Performance comparison for Expression Prediction task (sorted by RegHit auROC)

| Participant<br>(Lab-Submission) | Ref. Spearman corr. | Alt. Spearman corr. | RegHit auPRC | RegHit auROC |
| --- | --- | --- | --- | --- |
| 4 (EnsembleExpr) | **0.484936** | **0.470176** | **0.528288** | **0.807690** |
| 6-1 | 0.290971 | 0.399997 | 0.461099 | 0.786722 |
| 2-2 | 0.278613 | 0.262536 | 0.402448 | 0.777035 |
| 2-4 | 0.260072 | 0.245915 | 0.437067 | 0.774688 |
| 2-5 | 0.261064 | 0.245595 | 0.432639 | 0.772747 |
| 6-2 | 0.290971 | 0.399997 | 0.433406 | 0.771472 |
| 6-3 | 0.433043 | 0.399116 | 0.426697 | 0.767268 |
| 2-6 | 0.199247 | 0.171989 | 0.353660 | 0.728643 |
| 2-1 | 0.173587 | 0.169082 | 0.385369 | 0.723336 |
| 1-5 | 0.295519 | 0.272904 | 0.304145 | 0.719242 |
| 1-1 | 0.251873 | 0.248376 | 0.329400 | 0.716045 |
| 1-3 | 0.251027 | 0.248319 | 0.328427 | 0.713983 |
| 1-6 | 0.318642 | 0.300630 | 0.312914 | 0.713570 |
| 1-4 | 0.254598 | 0.236243 | 0.311588 | 0.709843 |
| 5-1 | 0.252023 | 0.223952 | 0.369462 | 0.693357 |
| 1-2 | 0.174655 | 0.168176 | 0.293471 | 0.683512 |
| 7 | 0.208036 | 0.194298 | 0.437487 | 0.670681 |
| 3 | 0.304933 | 0.236940 | 0.242830 | 0.652059 |
| 5-2 | 0.352951 | 0.353493 | 0.189516 | 0.578095 |
| 2-3 | 0.000766 | -0.002387 | 0.126747 | 0.513558 |

In the second part of the challenge, EnsembleExpr accurately predicted the change in expression (“LogSkew”) with a Spearman correlation much better than most of the other submissions which yielded close to zero (Table 2). Prioritizing variants that give rise to significant change of expression (“emVar”) is the hardest among all tasks. In this task, EnsembleExpr also demonstrated superior performance with a auROC of 0.655 and an auPRC of 0.452 (Table 2, Figure 2B).

**Table 2:** Performance comparison for Allele-specific Expression Prediction task (sorted by emVar auROC)

| Participant<br>(Lab-Submission) | LogSkew Spearman corr. | emVar auPRC | emVar auROC |
| --- | --- | --- | --- |
| 4 (EnsembleExpr) | **0.449760** | **0.452561** | **0.655261** |
| 5-1 | 0.333893 | 0.409730 | 0.626850 |
| Published state-of-the-art | Not Applicable | 0.389 | 0.589 |
| 5-2 | 0.342004 | 0.369083 | 0.577220 |
| 7 | 0.007343 | 0.431639 | 0.562854 |
| 6-1 | 0.217845 | 0.345064 | 0.561953 |
| 6-2 | 0.190123 | 0.354726 | 0.561776 |
| 1-3 | NaN* | 0.311243 | 0.556499 |
| 1-1 | NaN* | 0.305258 | 0.550820 |
| 1-2 | 0.030243 | 0.295886 | 0.550048 |
| 2-3 | -0.015476 | 0.303051 | 0.545206 |
| 1-5 | 0.056143 | 0.284863 | 0.541216 |
| 1-4 | 0.079049 | 0.293321 | 0.530856 |
| 3 | 0.030049 | 0.284356 | 0.511181 |
| 1-6 | 0.105376 | 0.286584 | 0.510103 |
| 2-2 | -0.007377 | 0.249473 | 0.479746 |
| 2-1 | -0.024347 | 0.234723 | 0.477301 |
| 2-5 | -0.023092 | 0.233144 | 0.472651 |
| 2-6 | -0.023092 | 0.233144 | 0.472651 |
| 2-4 | -0.023092 | 0.233144 | 0.472651 |

\*: every variant was assigned the same score, leading to incalculable Spearman correlations

EnsembleExpr also outperformed the state-of-the-art in the eQTL-prioritization literature. Recent work (Zhou and Troyanskaya, 2015) reported that a *L*_2_-regularized logistic regression trained on DeepSEA-derived features and evolutionary conservation scores achieved a performance that surpasses existing approaches. We generated the same features for the datasets in the eQTL challenge and trained the same regularized logistic regression model to predict “emVar” labels. While ranking third among all the submissions, this model achieved a performance inferior to that of EnsembleExpr (auROC=0.589, auPRC=0.389, Table 2). This comparison shows that EnsembleExpr not only excels among all the submitted methods, but outperforms the state-of-the-art in the literature.

Thus EnsembleExpr modeled the diversity of expression well and demonstrated unmatched capacity as a predictive model for eQTL prioritization. More importantly, the consistently high performance of EnsembleExpr across different tasks and evaluation metrics proves the robustness of the predictions.

### 4.2 Components of the ensemble provide complementary functional information

We benchmarked EnsembleExpr and each of the single models included in the ensemble to understand the major sources of improvement. Through ten-fold cross-validation, for each model we evaluated the median *R*^2^ when predicting log expression level (“Log2FC”) and the median auROC and auPRC when predicting significant expression (“RegHit”). We observed that with the DeepSEA-predicted functional features, including TF binding, histone marks and DNase hypersensitivity, we could already reach decent accuracy in both tasks (Table 3). However, models with only TF binding-based features from either deep learning (DeepBind) or k-mer based models (KSM) are much less satisfactory. But we did observe that incorporating DeepSEA with features from KSM and ChromHMM led to better performance, suggesting that these two models provided complementary information despite the comprehensiveness of the DeepSEA output.

**Table 3:** Performance of each component in the ensemble

| Features included | Task1-a | Task1-b |  |
| --- | --- | --- | --- |
| | $R^2$ | auROC | auPRC |
| Ensemble | **0.3976** | **0.8647** | **0.5830** |
| KSM+DeepSEA* | 0.3803 | 0.8515 | 0.5622 |
| KSM+DeepSEA+ChromHMM* | 0.3769 | 0.8462 | 0.5380 |
| DeepSEA* | 0.3728 | 0.8347 | 0.5391 |
| DeepBind* | 0.2508 | 0.8209 | 0.4438 |
| KSM | 0.2393 | 0.7943 | 0.3943 |

\*: models included in the ensemble

### 4.3 Accurate eQTL prioritization requires a comprehensive panel of functional features

We next sought to understand what sequence-derived functional features, among the hundreds we used, are most predictive of expression and eQTL status. Expression is regulated by sophisticated machinery where numerous regulators and epigenetic marks act in concert. To include a large enough panel of features, we investigated one of the LASSO regression models in the ensemble that was trained to predict expression (“Log2FC”) from sequence-derived prediction of DNase hypersensitivity, histone marks, transcription factor binding and chromatin state (Supplementary Table 1).

We first analyzed the sign of the coefficients in the LASSO model to understand the direction in which each feature affects the expression prediction. As expected, the model assigned large positive weights to DNase hypersensitivity, histone marks known to be associated with promoters (such as H3K4me3) and active functional elements (such as H3K27ac), and transcription initiators (such as IRF1) (Supplementary Table 2). The model also gave large negative weights to chromatin regulators known for repressive effects on transcription (such as EZH2) and histone marks predictive for gene bodies (such as H3K36me3). Interestingly, we observed that the LASSO model consistently assigned negative or close-to-zero coefficients to H3K4me1, which is known as strongly indicative for distal elements such as enhancers (Creyghton *et al.*, 2010). This observation persisted even when we retrained the model 10 times, and calculated the mean and 95% confidence interval of the coefficients (Supplementary Table 2). As DeepSEA does not predict H3K4me1 any less accurately than other marks (Supplementary Table 2 in Zhou and Troyanskaya (2015)), we speculate that this might reflect MPRA’s insensitivity to sequences that regulate gene expression in trans.

We next analyzed the importance of the features. By design, LASSO models impose sparsity and force the coefficients for non-important features to zero. However, the limitation of such *L*_1_-regularization based models is that when faced with a group of highly correlated features, as in our experiments, the model will only pick one of them. Thus to fully understand which features are important for expression prediction, instead of directly looking at the coefficients in the LASSO model, we retrained a Randomized Lasso model that performs “stability selection” (Meinshausen and Bühlmann, 2010) by resampling the train data and computing a LASSO model on each resampling. The more often a feature gets selected, the more important it is for the performance of the model. We observed a bi-modal distribution of feature importance (Supplementary Figure 1). Most of the 994 features are considered not very important, while a group of 60 features demonstrate great importance. These top 60 features are highly diverse, including histone marks predictive for enhancer/promoter/repressive regions, important transcription regulators and chromatin states predictions (Supplementary Table 3). This diversity of useful features suggests that a comprehensive functional annotation of the sequence, rather than one type or two, is essential for accurate expression prediction and eQTL prioritization. We also observed that while many of the important features are predicted for the same type of cell line as the one the MPRA experiment was performed on (lymphoblastoid cell lines), many features predicted for other cell lines, such as K562 and H1-hESC, also proved to be highly informative.

## 5 Discussion

In this work, we presented EnsembleExpr, an ensemble-based framework that predicts expression level from sequence and prioritizes sequence variants that exhibit allele-specific expression. We showed that EnsembleExpr achieved the best performance in both parts of the “eQTL-causal SNPs” challenge in the Fourth Critical Assessment of Genome Interpretation (CAGI4).

Each component of the EnsembleExpr provides useful yet complementary information, leading to a successful ensemble with performance surpassing any of the single ones. Through a systematic analysis of feature importance, we demonstrated that the features considered important for accurate prediction are highly diverse, ranging from chromatin state and histone marks to transcription factor binding.

In this framework, most of the features we used, except the chromatin state labels from ChromHMM, are obtained from sequence-based computational models that can provide allele-specific predictions. This enables precise characterization of how a single-base change affects expression levels, which we consider crucial for any model aiming to interpret sequence variants.

With the capacity to accurately predict sequence variants with significant allele-specific expression, we expect EnsembleExpr to serve as an important resource to pinpoint mutations causal for complex traits and diseases and help understand the pathogenic pathways. We make EnsembleExpr openly available at http://ensembleexpr.csail.mit.edu for researchers to utilize freely for downstream analysis.

## Acknowledgments

We are grateful to the organizers of the Fourth Critical Assessment of Genome Interpretation for coordinating and hosting the open challenge. We appreciate Ryan Tewhey and Pardis Sabeti from the Broad Institute for providing the MPRA data in the “eQTL-causal SNPs” challenge. We thank Kevin Tian for the help in feature preparation. We thank other members of the Gifford Lab for constructive discussions and feedback. We acknowledge funding from the National Institutes of Health under grants R01HG008363 and U01HG007037 to D.K.G. and an equipment grant from NVIDIA.

